# Pyruvate dehydrogenase dependent metabolic program affects oligodendrocyte maturation and remyelination

**DOI:** 10.1101/2023.03.19.533192

**Authors:** M Sajjad, Insha Zahoor, Faraz Rashid, Ramandeep Rattan, Shailendra Giri

## Abstract

The metabolic need of the premature oligodendrocytes (Pre-OLs) and mature oligodendrocytes (OLs) are distinct. The metabolic control of oligodendrocyte maturation is not fully understood. Here we show that the terminal maturation and higher mitochondrial respiration in the oligodendrocyte is an integrated process controlled through pyruvate dehydrogenase (Pdh). Combined bioenergetics and metabolic studies show that mature oligodendrocytes show elevated TCA cycle activity than the premature oligodendrocytes. Our signaling studies show that the increased TCA cycle activity is mediated by the activation of Pdh due to inhibition of pyruvate dehydrogenases isoform-1 (Pdhk1) that phosphorylates and inhibits Pdh. Accordingly, when Pdhk1 is directly expressed in the premature oligodendrocytes, they fail to mature. While Pdh converts pyruvate into the acetyl-CoA by its oxidative decarboxylation, our study shows that Pdh also activates a unique molecular switch required for oligodendrocyte maturation by acetylating the bHLH family transcription factor Olig1. Pdh inhibition via Pdhk1 blocks the Olig1-acetylation and hence, oligodendrocyte maturation. Using the cuprizone model of demyelination, we show that Pdh is deactivated during the demyelination phase, which is reversed in the remyelination phase upon cuprizone withdrawal. In addition, Pdh activity status correlates with the Olig1-acetylation status. Hence, the Pdh metabolic node activation allows a robust mitochondrial respiration and activation of a molecular program necessary for the terminal maturation of oligodendrocytes. Our findings open a new dialogue in the developmental biology that links cellular development and metabolism. These findings have far-reaching implications for the development of therapies for a variety of demyelinating disorders including multiple sclerosis.

## Introduction

Oligodendrogenesis is a multistep process in which oligodendrocyte progenitor cells (OPCs) first differentiate into the oligodendrocyte linage and then undergo terminal maturation into the myelinating oligodendrocytes [1]. Oligodendrogenesis is critical to the functioning of CNS, given oligodendrocytes myelinate neurons which maintains the tissue integrity as well as speed of the nerve transmission [1-3]. Hence, oligodendrogenesis is central to the behavior and cognitive ability of an organism [4,2,5]. Accordingly, disorders affecting this process can impair the cognitive abnormalities of an organism to varying degrees [6-8]. One such example is multiple sclerosis (MS), where in the immune system attacks the myelin leading to its degradation along with the stalling the oligodendrogenesis at multiple stages. These events denude the axons and slow the speed of conduction thus affecting the cognitive and motor abilities of the affected patients [9-14].

The current wealth of research in the oligodendrogenesis field has identified specific molecular players dictating the early fate attainment of OPCs [15-18] and the transcriptional programs regulating oligodendrogenesis [17], however understanding of the terminal maturation process of oligodendrocytes is still developing. This becomes especially important in the context of MS, since recent investigations using the patient CNS tissues has revealed that while the early OPC differentiation is not grossly affected, the disease stalls the terminal maturation of the premature oligodendrocytes [19]. Hence, understanding the process of terminal maturation of oligodendrocytes is necessary to advance the CNS biology as well as for the pursuit of effective repair therapies for MS.

The biology of oligodendrocyte terminal maturation is still not well understood. It is worthwhile to note that terminal maturation of the oligodendrocytes is a metabolically active process. Recently, the central role of cellular metabolism in the terminal maturation of the oligodendrocytes has been emerging [20-23]. A recent study demonstrated that ATP generation via TCA cycle was a critical determinant of the OPC maturation into Pre-oligodendrocytes and mature oligodendrocytes which display significantly higher mitochondrial respiration than OPCs [24]. Another study using in-depth metabolism tracing using the ^13^C-acetate labeling showed that acetate is efficiently converted to acetyl-CoA in the mature oligodendrocytes and fuels the TCA cycle [25]. Studies have also shown rapid increase of mitochondrial genes during oligodendrocyte differentiation and its inhibition by its repression [26]. Further evidence in this direction comes from the work showing robust metabolism in oligodendrocytes post injury supporting the latter notion [27] apart from direct evidence of the need of mitochondrial respiration during oligodendrocyte maturation [26]. These findings are further strengthened by the fact that metabolic enhancement of the aged OPCs leads to their better-maturation [28]. Despite of these studies, we have not been able to pin point how integration of metabolic programming and signaling and metabolic processing underlying increased mitochondrial respiration during oligodendrocyte maturation especially during the transition of the premature oligodendrocytes into the mature oligodendrocytes remains unknown. Accordingly, using a combined approach of the signal transduction and metabolomics, we report pyruvate-dehydrogenase (Pdh) as a critical node that regulates oligodendrocyte maturation from the premature oligodendrocytes. More specifically, the Pdh is activated during transition from the premature to mature oligodendrocytes and provides a dual edge by allowing increased energy production via TCA cycle and modulation of cellular signal apparatus via acetylation during the maturation of the oligodendrocytes.

## Results

### Mitochondrial respiration is elevated during the pre-OL maturation into the Ols

To profile the bioenergetics signatures during the maturation of pre-OLs into OLs, we analyzed rat cultured OPC differentiated into pre-OLs and OLs (**Fig.1A**) using the Seahorse bioenergetics analyzer XF^e^96 ™ which reveals bioenergetics state of live cells in terms of glycolysis (extracellular acidification rate; ECAR) and mitochondrial respiration (oxygen consumption rate; OCR). We observe that OLs utilize significantly higher OCR than pre-OLs (**Fig. 1B & C**). Previous study showed pre-OLs to exhibit significantly higher OCR than OPCs comparing OPCs with pre-OLs indicated a striking difference in mitochondrial respiration, with pre-OLs possessing higher OCR than OPCs [24]. Together, these findings indicate that the process of OL differentiation and maturation is accompanied by an increased mitochondrial respiration. Since, glycolysis and TCA cycle are interconnected, we also analyzed the glycolytic rate under similar conditions by measuring the ECAR between pre-OLs and OLs, which showed a decreased ECAR in OLs compared to Pre-OLs (**Fig. 1B & C)**. Our bioenergetics studies also revealed net higher energy state of the OLs vs the pre-OLs (**Fig. 1D)**. To interrogate these findings more intensively, we labeled the pre-OLs and OLs with U-13C glucose and performed the LC/MS analysis of the U-C13 enrichment in citrate to monitor flux into the TCA cycle. Our data shows that OLs had higher U-13C citrate enrichment (**Fig. 1E)** when compared to the pre-OLs. This data suggests that OLs show higher OCR due to increased utilization of the glucose in the TCA pathway. Together these findings imply that during transition from Pre-OLs to OLs, glucose utilization shifts toward the mitochondrial respiration.

**Figure 1:**
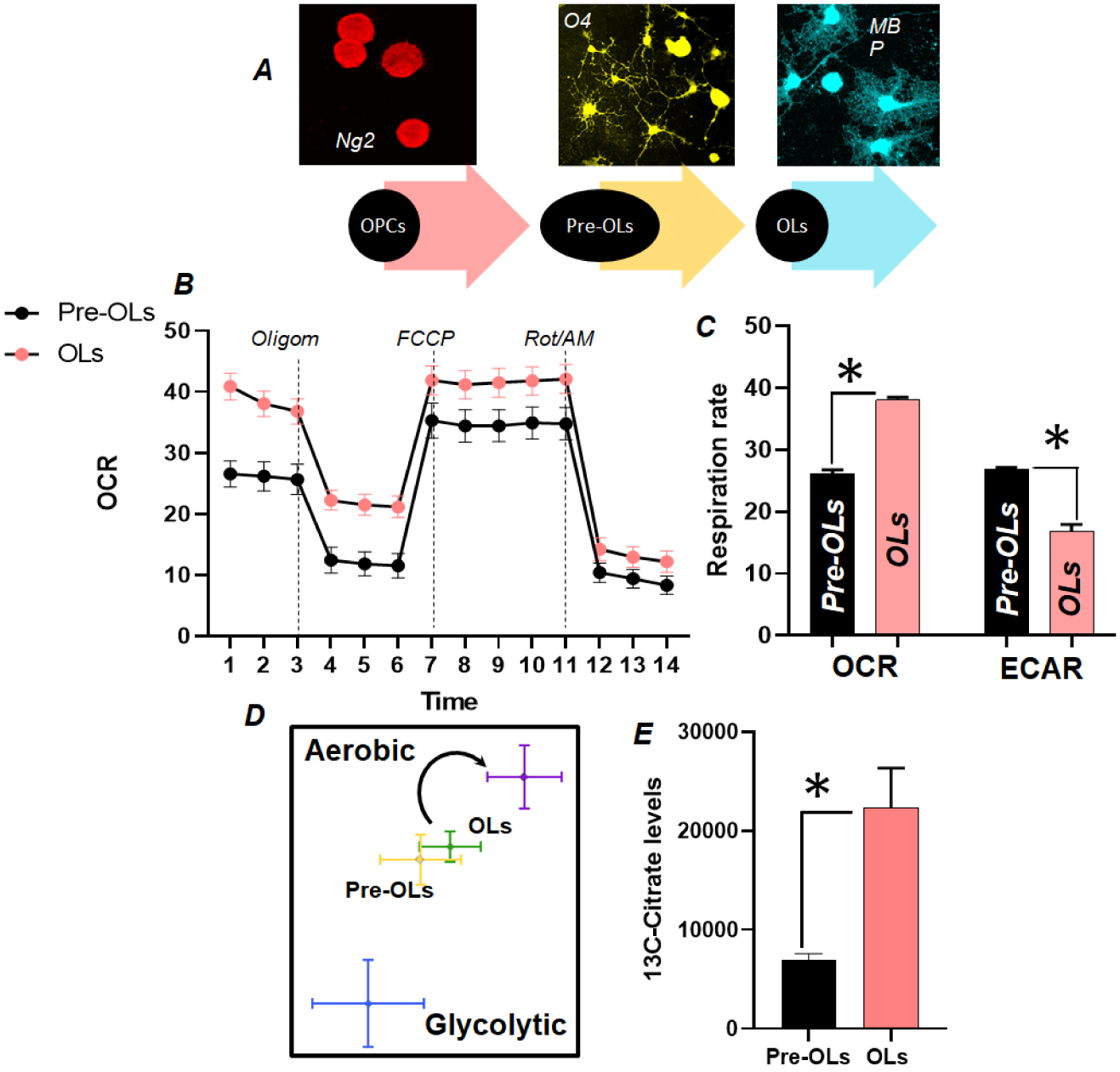
OLs show elevated mitochondrial respiration. **A-** Rat OPCs were cultured from cortical cultures. The OPCs were initially propagated as Ng2 expressing oligospheres and subsequently differentiated into the O4 expressing Pre-OLs and finally MBP expressing OLs. **B**- Immature O4+ and mature MBP+ oligodendrocytes were plated in the Seahorse cell plate from 3-days in order to have 2 different differentiation stages. Bioenergetics of the cells was conducted using extracellular flux analyzer. The cells were treated with the Oligomycin, FCCP and Rapamycin/Antimycin in different stages of the experiment. The oxygen consumption rate (OCR) as a measure of mitochondrial respiration was expressed as pMol/min and glycolytic acidification rate or ECAR was expressed as mPH/min. **C**- Graph showing the elevated OCR (p<0.01, n=18 per cell type) and reduced ECAR (p<0.01, n=18/cell type) in the OLs vs the Pre-OLs. **D-** Energy map of the Pre-OLs and OLs showing the higher Energy status and shifting of the respiration from the glycolysis to the oxidative phosphorylation in the OLs vs the Pre-OLs (black arrow). **E-** Pre-OLs and OLs were treated with U 13C glucose and LC/MS analysis was conducted to study the enrichment in the citrate from the TCA cycle (p<0.05, n=3/cell type). All the data expressed are Mean ± S.E.M

### Pdh-function is upregulated during Pre-oligodendrocyte (pre-OL) maturation into the Ols

Pyruvate produced from glycolysis is a switch point between lactate and mitochondrial respiration. Pyruvate enters the mitochondria and is decarboxylated into acetyl-CoA, which fuels both TCA cycle and *de-novo* fatty acid production [29]. The enzyme pyruvate dehydrogenase (Pdh) catalyzes the oxidative decarboxylation of pyruvate into acetyl-CoA. We next examined the Pdh activity during pre-OLs maturation into OLs and found a significantly increased Pdh activity in OLs compared to Pre-OLs (**Fig. 2A**), suggesting that higher mitochondrial respiration in OLs is possibly due to increased Pdh activity. Pdh is a multi-enzyme complex, coupling the activity of three component enzymes including pyruvate dehydrogenase (Pdh, E1), dihydrolipoamide acetyltransferase (E2) and dihydrolipoamide dehydrogenase (E3) in the oxidative decarboxylation of pyruvate to acetyl-CoA, linking glycolysis and TCA cycle. Specific inhibitory phosphorylation by a group of four evolutionary conserved Pdh kinases (Pdhk1-4) on the E1α subunit of the heterotetramer inhibit the activity of Pdh. In order to gain a mechanistic understanding of the elevated Pdh function during pre-OL maturation allowing higher oxygen consumption in OLs, we examined the expression of various isoforms of Pdh kinases and found a significant decrease in the expression of Pdhk1 isoform of the Pdh kinases (**Fig. 2B**), with very little or no effect on the expression of Pdhk2, pdhk3 and Pdhk4 (**Fig. 2B**). This observation is further supported by immunocytochemical analysis of cultured cells exhibiting lower expression of Pdhk1 in mature OLs expressing MBP compared to pre-OLs expressing O4. Consistent with the loss of Pdhk1, we observed a decrease in phosphorylation at E1-α subunit of the Pdh at Ser-293 [30] during this transition, which demonstrates that Pdh is activated during OL maturation possibly due to inhibition of Pdhk1. Overall, our data indicates that OLs adapt to an increased mitochondrial respiration possibly due to increase in Pdh activity, maybe due to inhibition of Pdhk1, which enables increased conversion of pyruvate to acetyl-CoA and increased fueling of TCA cycle.

**Figure 2:**
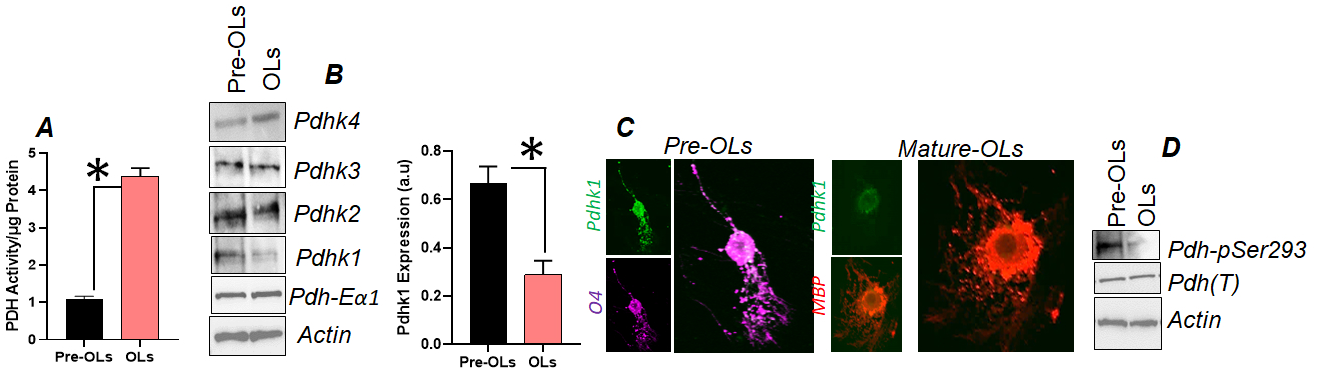
Pdh activity increases through Pdhk modulation during Pre-OL maturation into the OLs. **A-** Rat immature oligodendrocytes and mature oligodendrocytes were cultured from the cortices and cell extracts were prepared and subjected to the colorimetric Pdh activity test which showed higher Pdh activity in the mature oligodendrocytes (p<0.05, n=3/per cell type).**B-** immunoblotting was conducted on the cellular lysates from the immature and mature oligodendrocytes for Pdhk1, Pdhk2, Pdhk3, Pdhk4 and Pdh total (T) in the immature and the mature oligodendrocytes. Pdhk1 expression in the Pre-OLs and OLs was quantified and normalized to the actin levels in the samples (p<0.05, n=3). **C-**Laser scanning confocal microcopy for Pdhk1 (green) expression in the O4 expressing (magenta-pseudocolor) immature oligodendrocytes and MBP expressing (red) mature oligodendrocytes. Please notice the reduced Pdhk1 expression in the MBP expressing mature oligodendrocyte. **D-**Immunoblotting for total and phosphorylated Pdh at Ser-293 in the Pre-OLs and OLs. All the data expressed are Mean ± S.E.M

### Pdh deactivation by Pdhk1 directly inhibits Pre-OL maturation into the OLs

After identifying that Pdh activity is elevated during Pre-OL maturation due to inhibition of Pdhk1, we examined if overexpression of Pdhk1 can directly impact the maturation of Pre-OLs into OLs. We transfected flag-expressing human Pdhk1 (*h*Pdhk1) (Addgene: 20564) in pre-OLs following partial differentiation from OPCs and monitored the maturation of pre-OLs into OLs. Anti-flag immunocytochemistry was used to evaluate the transfection of flag-*h*Pdhk1 in O4 expressing pre-OLs (**Fig. 3A**). Pdh deactivation by overexpressed Pdhk1 was further investigated by-immunoblotting against phospho-Pdh (Ser-293) antibodies. Overexpression of flag-*h*Pdhk1 transfected pre-OLs showed a decrease in Ser-293-phosphorylated Pdh indicating inhibition of Pdh activity (**Fig. 3A**), which was confirmed by assessing its enzymatic activity under similar experimental conditions (**Fig. 3B**). To examine the effect of Pdh deactivation on the maturation of pre-OLs into OLs, we transfected pre-OLs either with a scrambled vector (scram) or flag-*h*Pdhk1. MBP-expressing myelinating OLs were counted in both groups. Overexpression of flag-*h*Pdhk1 in pre-OLs, caused a significant reduction in the number of pre-OLs maturing into OLs compared to scram-transacted pre-OLs (**Fig. 3C**). Furthermore, we observed less membrane expression in the differentiated OLs of the Flag-hPdhk1 transfected group than the scram transfected group **(Fig.3C**). Together, our findings show that Pdh deactivation by Pdhk1 can impede pre-OL maturation, suggesting that Pdh driven oxidative phosphorylation is critical in the maturation of premature oligodendrocytes into myelinating oligodendrocytes.

**Figure 3:**
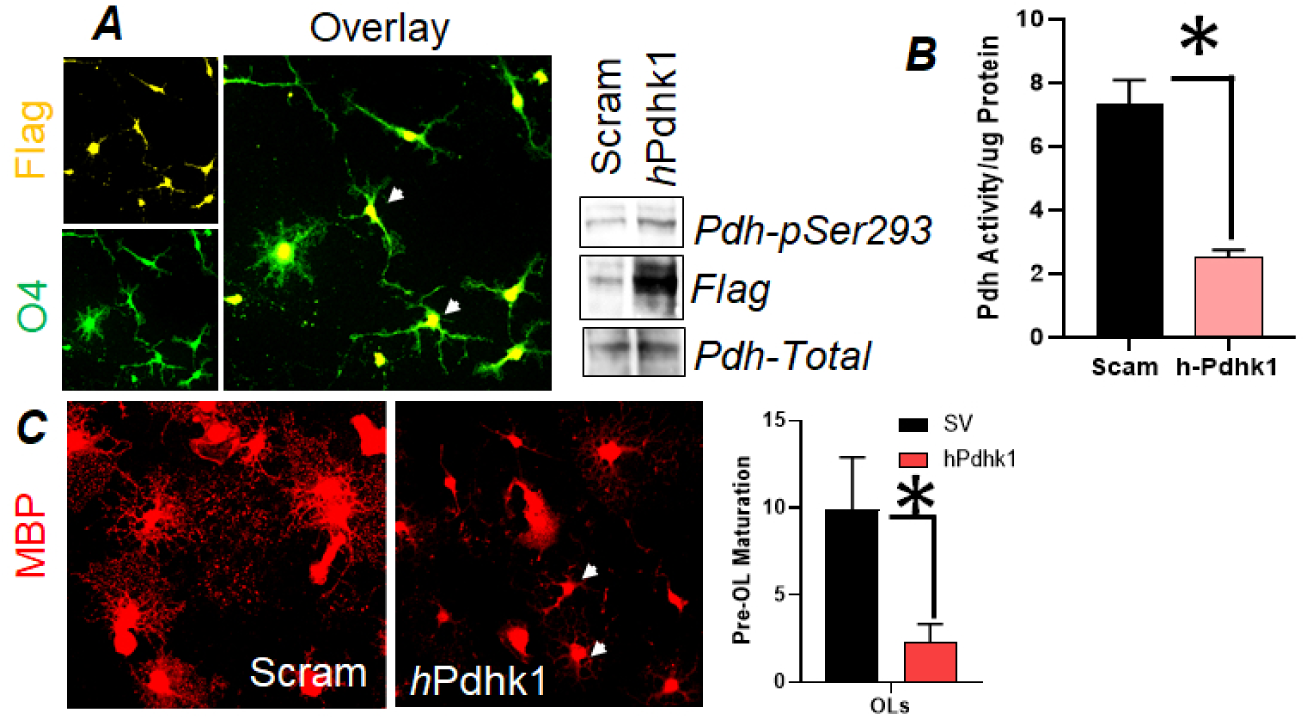
Pdhk1 mediated Pdh deactivation arrests Pre-OL maturation into the OLs. **A-** Rat OPCs were partially differentiated into the Pre-OLs and transfected with either scrambled-hPdhk1 (SV) or intact Flag-*h*Pdhk1 vector (*h*Pdhk1) for 12h. **A-**Laser scanning confocal microscopy was performed to monitor the Flag- *h*Pdhk1 (pseudo-yellow) expression in the O4+ Pre-OLs (green) at 72h later after fixing the cells. Images clearly showed Flag expression in the O4+ immature OLs. Sister cultures were immunoblotted for the for the Pdh phosphorylation at the Ser-293 meaning its deactivation while using Flag from the expression vector and total Pdh as the experimental controls. Scrambled *h*Pdhk1 (SV) or intact Flag-*h*Pdhk1 vector (*h*Pdhk1) were transfected in the rat Pre-OLs. **B-** The cell extracts were probe for Pdh activity (p<0.01, n=3/cell type) at 72h post transfection. **C-** Scrambled *h*Pdhk1 (SV) or intact Flag-*h*Pdhk1 vector (*h*Pdhk1) were transfected in the rat Pre-OLs and were allowed to differentiate in the standard differentiating conditions and laser scanning confocal microscopy was performed for the MBP expressing mature OLs. The MBP expressing OLs were counted from multiple regions of interest from both the groups (p<0.01, n=4 ROIs per group). All the data expressed in the results are Mean ± S.E.M.

### Pdh inhibition by Pdhk1 blunts the acetylation and nuclear to cytoplasmic shuttling of Olig1 during the Pre-OL maturation into the OLs

Together from the bioenergetics signature, signal and metabolic studies, it’s clear that elevated TCA function in the in the OLs is an outcome of the elevated Pdh activity. Given Pdh activity decarboxylates the pyruvate into the acetyl-CoA that delivers critical acetyl groups to the TCA via the citrate [31], we estimated the acetyl-CoA levels in the pre-OLs and the OLs. We observed a significant rise in the acetyl-CoA levels in the OLs compared to pre-OLs (**Fig. 4B** which confirms that during the pre-OL maturation, the elevated Pdh activity drives more pyruvate into acetyl-CoA production. Acetyl-CoA, not only fuels the TCA but also acts as substrate for the process of acetylation [32-35]. It’s known that acetylation of the bHLH family transcription factor Olig1 is critical to its nuclear to cytoplasmic shuttling and maturation of the OLs [36]. Upon acetylation, the Olig1 relocates from the nucleus to the cytoplasm and enables the differentiation of pre-OLs into the OLs [36]. Previous studies have demonstrated higher Olig1 acetylation in the OLs compared to pre-OLs. [36]. We postulated that increased Pdh activity led to generation of increased acetyl-CoA from pyruvate, which maybe critical for Olig1 acetylation and nuclear to the cytoplasmic translocation and hence OL maturation. To confirm this, we used confocal imaging to track the subcellular localization of Olig1 during the transition from pre-OLs to OLs. We observed higher cytoplasmic localization of Olig1 in OLs in contrast to the pre-OLs (**Fig. 4A)**. To confirm further, we differentiated the pre-OLs with or without the ectopic expression of *h*Pdhk1 to deactivate the Pdh for 3 days and monitored the cytoplasmic expression of Olig1. We observed elevation in the cytoplasmic Olig1 in the scrambled vector transfected differentiated pre-OLs as before (**Fig.4C**), while the *h*Pdhk1 expressing pre-OLs did not show any significant increase in the cytoplasmic Olig1. These results also agree with the above findings in which *h*Pdhk1 expression blunts the generation of MBP+ mature OLs from the pre-OLs. We also monitored the Olig1 acetylation in the vector control and the *h*Pdhk1 expressed pre-OLs by co-immunoprecipitation followed by the immunoblotting which revealed reduced Olig1-acetylation in the *h*Pdhk1 expressing pre-OLs (**Fig.4E**) confirming that expression of *h*Pdhk1 by deactivation the Pdh reduces acetyl-CoA delivery to the Olig1 apart from reducing the vital energy pathway and thus blunts their maturation.

**Figure 4:**
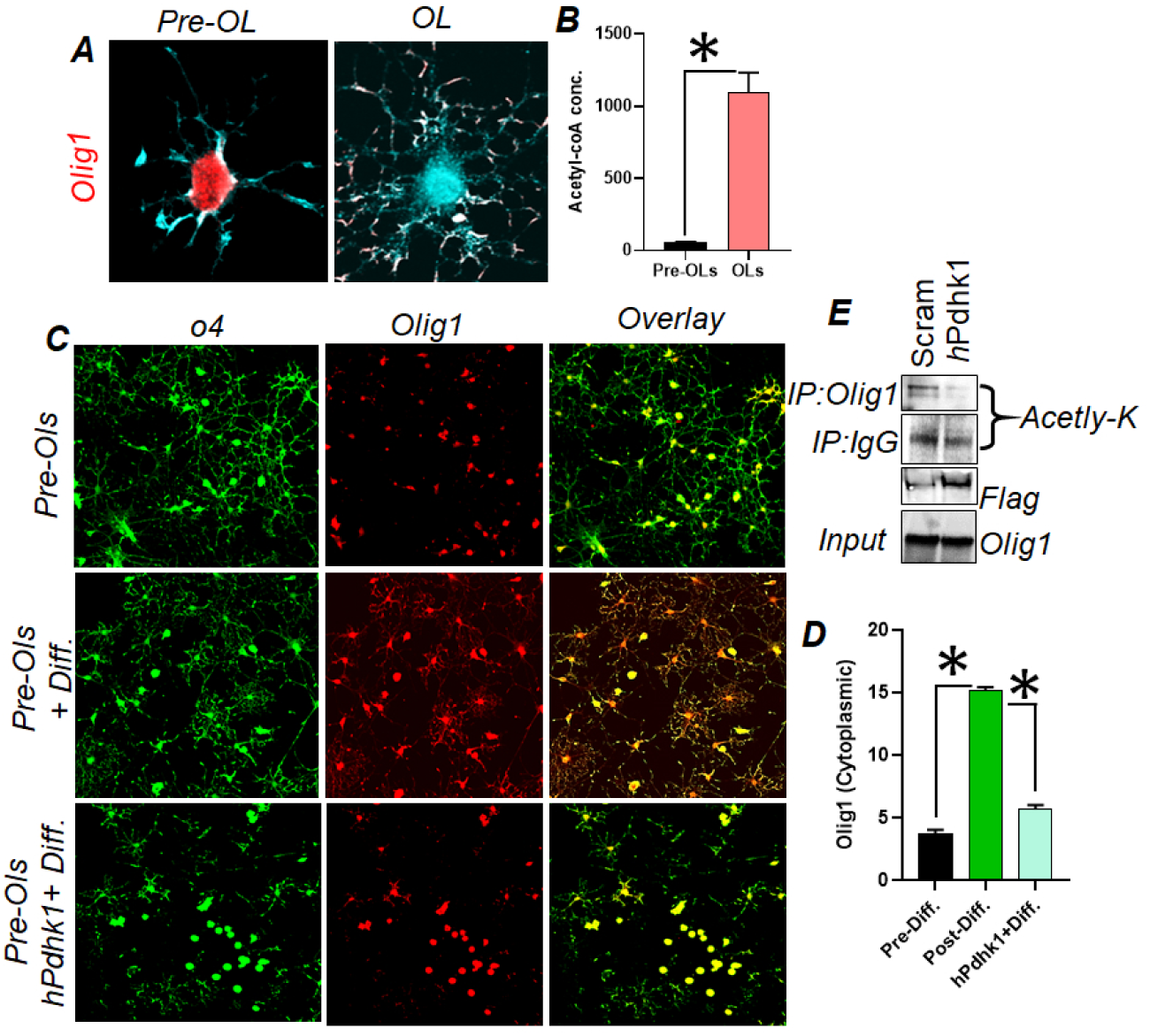
Pdhk1 through acetyl-CoA production controls the Olig1 mediated Pre-OL maturation. **A-** OPCs were isolated from the Rat pups and differentiated into Pre-OLs (O4-Cyan pseudo color) and OLs (MBP-Cyan, pseudo color). Immunocytochemistry followed by the laser scanning confocal microscopy was conducted for Olig1 expression (Red) in the Pre-OLs and OLs (white arrows pointing to the Olig1 expression in the Pre-OLs and the OLs). **B-** LC/MS analysis of the Pre-OLs and OLs for the total acetyl-CoA in the Pre-OLs and the OLs (p<0.01, n=3/cell group). **C-** Rat OPCs were generated as described and differentiated under the standard differentiation conditions with or without the transfection for *h*Pdhk1 and immunocytochemistry followed by the laser scanning confocal microscopy was performed for O4 (green) and Olig1 (red) to check the nuclear to cytoplasmic shifts of Olig1 and its impact by the *h*Pdhk1 (white arrows show cytoplasmic expression of Olig1). **D-** Cytoplasmic expression was quantified by the intensity measurements from multiple cells in the group using Image-J (Pre-Diff vs. the Post diff., p<0.01, n=20 cells, Post diff., vs Post Diff+ *h*Pdhk1, p<0.01, n=20 cells). **E-** Pre-OLs were either transfected with the scrambled hPdhk1 or Flag- *h*Pdhk1 for 12h and the cell extracts at 72h were co-immune precipitated with either nonspecific IgG or Olig1 specific antibody. The co-immune precipitates were further probed by immunoblotting for pan-acetylated lysine antibody. All the data expressed in the results are Mean ± S.E.M.

### Pdh activity determines Olig1 acetylation status and the outcomes of remyelination in the cuprizone model

We used the cuprizone-chemical induced demyelinating mouse model, which allows to track demyelination and remyelination via cuprizone dietary modification [37,38]. Previous studies have found an elevated level of Pdhk1 during the demyelination phase of this model [39]. We first monitored the Pdh activity at the base line, week-1 and 4 after starting cuprizone diet and week-3 post withdrawal of the cuprizone. The results showed a decrease in Pdh activity at week one, which progressed to further decrease at week four (**Fig. 5A**) post-cuprizone diet. However, we observed an increase in Pdh activity at week 3 after cuprizone diet withdrawal, which corresponds to the remyelination phase in this model (**Fig. 5A, p<**). We also found that the Pdhk1 expression and Pdh Ser-293 phosphorylation pattern also correlated with the Pdh activity status as observed above (**Fig. 5B**). We observed Olig1 acetylation at the same time points and found that Pdhk1 mediated Pdh deactivation leads to the Olig1 de-acetylation in the demyelination phase (week 1 and week-4) and rebounds with the resurgence of Pdh activity in the remyelination phase, week 3 post cuprizone withdrawal (**Fig. 5C**). As a further layer of evidence, we performed the flourmylein stain in addition to the immunofluorescence for Olig1 to distinguish it’s nuclear to cytoplasmic shifts, as well as the co-immunostain for pPdhk1 in O4 expressing pre-OLs. As per the imaging results, Olig1 is primarily sequestered in the nucleus during demyelination and begins to display cytoplasmic expression during the remyelination phase (**Fig. 5D**). Moreover, O4-expressing pre-OLs have higher phospho-Pdh (Ser-293) levels during demyelination and lower levels during remyelination (**Fig. 5E**). Taken together, our results suggest that Pdh activity is important for OL maturation both *in vivo* and *in vitro*.

**Figure 5:**
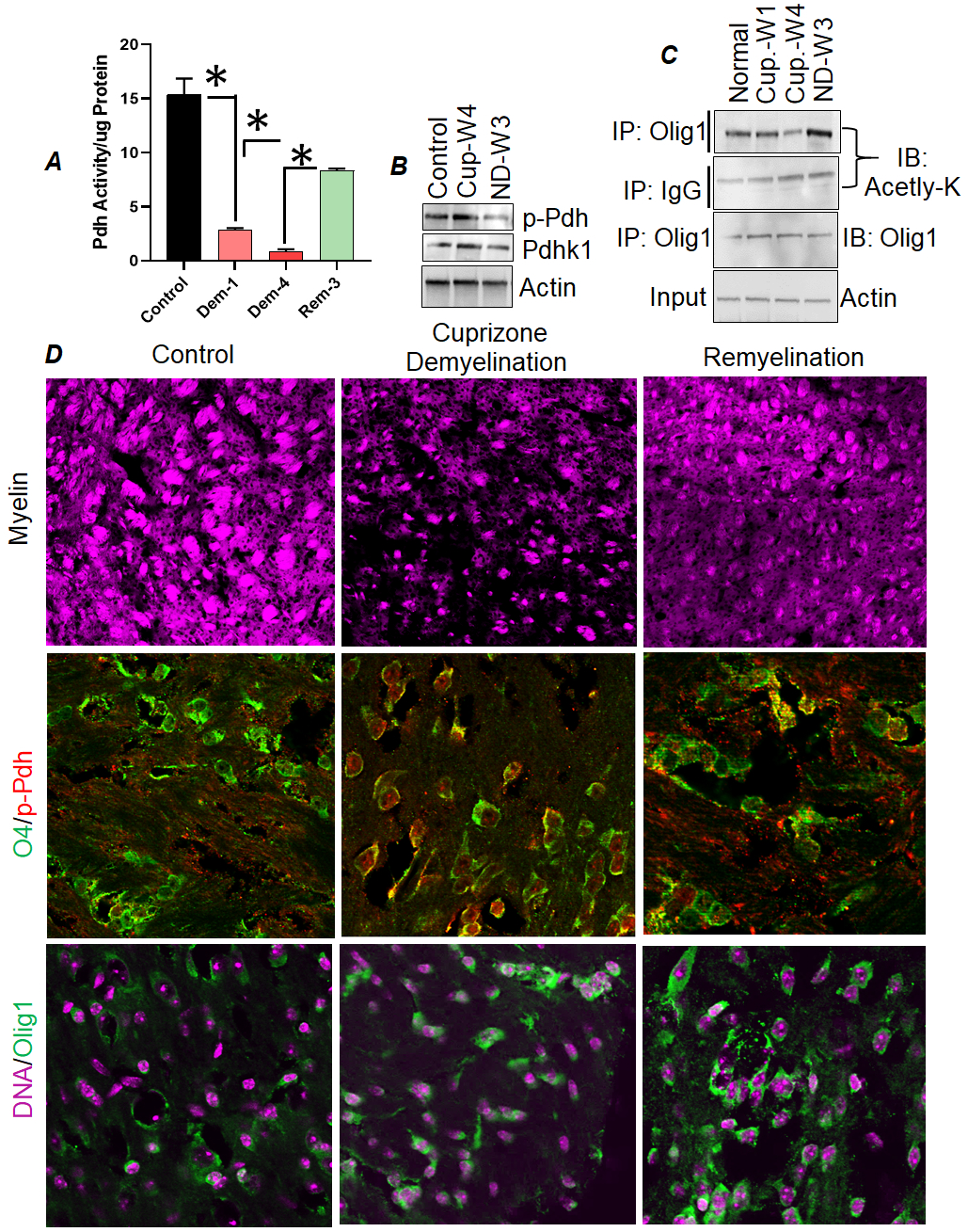
Pdh activity regulates olig1 acetylation and remyelination. **A-** Pdh activity was probed in the control mice, cuprizone (0.25%) demyelinated mice at 1 (Dem-1), 4 weeks post cuprizone diet (Dem-4) and 3 weeks post induction of the normal diet after cuprizone treatment (Rem-3) (n=3/time point in all the groups, control vs. dem-1, p<0.01, dem-1 vs dem-4, ns and Dem-1 vs Rem-3, p<0.01 and Dem-4 vs Rem-3, p<0.01). **B**- Immunoblotting for Pdhk1 and p-Pdhk1in the control mice, Dem-4 and Rem-3. **C**- Co-immunoprecipitation was carried using Olig1 antibody and a nonspecific isotype matched IgG antibody in the Control, Dem-1, Dem-4 and Rem-3 mice in the white matter followed by the immunoblotting for pan Acetyl-lysine (K) antibody. Internal controls employed were input precipitated Olig1 and actin. **D-**Confocal laser scanning imaging of the white matter of the control mice, Dem-4 and Rem-3 mice. The myelin was stained with the coronal sections with flour myelin, premature oligodendrocytes were labeled with O4 (green and co-expression was checked for phosphor-Pdh (red) (white show the p-Pdh in the Pre-OLs). Similarly Olig1 (green) expression was imaged with nuclear contrast by DAPI (magenta, pseudo color) (White arrows show the nuclear expression of Olig1). All the data expressed in the results are Mean ± S.E.M

## Experimental Methods

### Chemical and Reagents

Added in the methods at appropriate places with catalog information.

### Animal Ethical Compliance at HFH

All the mouse work presented in this manuscript was authorized the IACUC committee at the Henry Ford Health. HFH is compliant with the national guidelines under the assurance number D16-00090. The mice used the experiments were mixed gender B6 mice of 8-10 weeks of age housed in the HFH vivarium at controlled 12:12 light and dark cycle with standard chow and *ad libitum*.

### Rat OPC differentiation into immature and mature oligodendrocytes

Post-natal rat pups at day-2 (P2) were sacrificed and mixed glia were prepared from the subventricular (SVZ) and cortical regions of the brain in high glucose DMEM supplemented with 10% F.B.S as per our already published methods [40] in T75 flasks with standard cell culture treatment. The medium was changed every 2-3 days. After about 7-9 days the T75 flasks containing the mixed glial cultures were shaken in an orbital shaker at 180 r.p.m for 18h at 37C. The cells were spun at 2000 r.p.m for 7min. and allowed to attach on a petri dish for about 2-5 min to let the non OPC cells adhere allowing to retain only free floating enriched OPCs. Following this procedure, cells were plated as per the needed of the experiment in 24 wells, 6-wells or 10 cm dishes in DMEM: F12 supplemented with PGDFα (20 ng/mL,Peprotech:100-13A), B27 (1%, Gibco:A3582801) and 10 ng/mL of FGF2, Peprotech:100-18B for 2-4 days. The differentiation was allowed to proceed for 4 days at which most of the cells are O4+. Further differentiation of the O4+ cells under T3 (100nM) stimulation converts O4+ Pre-OLs into the mature oligodendrocytes expressing MBP termed at OLs. The purity of the cultures was judged by the immunocytochemistry for O4. These revealed that our cultures are 90-95% pure with a possibility of 2-4% cells belonging to other identity.

### ^13^C-labeling, cell-treatments and transfections of the Pre-OLs

Depending upon the need of the experiment, the cells were treated in either 24 well plates, 6 well plates or the 6 well plates. In order to generate Pre-OLs and OLs, most of the cases the OPCs were differentiated into the premature oligodendrocytes as described above. For the transfection with human Pdhk1 clone (*h*Pdhk1; Addgene: 20564). Pre-OLs after partial differentiation for 4 days from the OPCs, were used for transfection using the Polyplus Jeprime reagent, a generous gift from the manufacturer (Catalog: 101000027). For a typical 6-welll dish, about 3uL/well of the *h*Pdhk1 construct were mixed with the transfection buffer (100uL) and mixed vigorously for 3-5 minutes. Following this, the transfection reagent (2uL) was added to the mixture and allowed to stand for the lipid-DNA complex formation at the room temperature with occasional mixing using a vortex shaker. The mixture was directly applied on the serum free premature oligodendrocyte cultures and the medium was shaken to allow homogenous mixing. The transfection was allowed to occur for 12h following which the medium was changed. For the experiments involving OL quantification after hPdhk1 transfection, the Pre-OLs were further differentiated for additional 4-5 days in DMEM: F12 containing the T3 (100nM). In case of the Olig1 studies in the Pre-OLs, the cells were maintained in the DMEM: F12 medium with PGDFα (20 ng/mL, Peprotech: 100-13A) after the *h*Pdhk1 transfection without T3. For ^13^C labeling of the Pre-OLs and OLs, the cells were mixed with 1uL/mL of the medium from the stock of U^13^C6-glucose (10mM, Sigma) for 5 hours. The cells were immediately scraped on ice in the chilled 80% methanol for the metabolic profiling for 13C Citrate enrichment and acetyl-CoA studies.

### Real time bioenergetics evaluation of the Pre-OLs and OLs by the XF6-Extracellular flux analyzer

Rat OPCs were cultured from the mixed glial cultures as described above. The barcoded 96-well Seahorse cell plates (Agilent: 101085-004) proved by the manufacturer (Agilent Technologies, U.S.A) were coated with PDL for 12-14h in dark at 37C. After 12-14hrs, the PDL was removed and the wells were washed with P.B.S for three time to wash out the excessive PDL. The cells were plated in the wells at a seeding density of ≥ 80,000 cells per well in DMEM: F12 (Gibco: 11320033). For terminal maturation of the OPCs, medium was supplemented with PGDFα (20ng/mL) and B27 (1%) for 2-3 days followed by per day treatment of T3 (100ng/mL). For the partial differentiation of the OPCs, the cells were first, maintained in FGF2 (10ng/mL) for 2-3 days and then allowed to differentiate using the above cocktail. This allowed us to have both premature and mature oligodendrocytes in a same plate for flux analysis. Before a day of the analysis, analytic plate was soaked in water and allowed to stand at 37C in a humidified chamber. On the analyses day, the water from the analytic plate was shifted to the manufacturer provided equilibration buffer (Agilent: 100840-000) for 2hrs before analysis. The medium from the cell plates was rapidly aspirated from the wells and new Seahorse analysis medium (DMEM without phenol red, Agilant:103576-100) at p.H-7.4 with 250uL of 1mMglucose, 250uL of 1mM glutamine and 250uL of 1mM pyruvate) was infused at the same time. The cells were monitored by two investigators for integrity before actual assay and maintained at 37C in a humidified chamber. The analytic plate was injected with Oligomycin (2uM), FCCP (500nM), Rotenone (1uM) and Antimycin (1uM) for the OCR (mitochondrial respiration) and Glucose (????) and 2DG (???) for extra cellular acidification rate (ECAR) as the measure of glycolysis. Analytic plate was inserted into the extracellular flux analyzer and allowed to optimize oxygen and p.H as described by the manufacturer. After that cells plate was replaced by the analytic plate for hands free operation for OCR and ECAR quantification. The OCR is expressed as pMol/min. and ECAR is expressed as mpH/min.

### Metabolite extraction, mass spectrometry and data analysis for LC/MS

Approximately, Two-million cells were harvested and quenched before metabolite extraction using the following procedure. Cells were rapidly rinsed by gently dispensing 5 mL of 37 °C deionized water to the cell surface. The plate was rocked briefly and aspirated, and quenched by directly adding 15 mL of Liquid Nitrogen to the dish. A volume of 100 µL water (0.2% Formic acid) was added and vortexed to mix, and 10 µL of ISTD dilution mix was added. Subsequently, 400 µL of acetonitrile (0.2% formic acid) was added and vortex to mix. Three cycles of extraction were carried out by vortexing for one minute followed by sonicating for 1 min (each). After this, the samples were centrifuged at 14,000 rpm for 15 min at 40C. Supernatant were loaded into pre-conditioned Phenomenex, Strata XL-100, 30mg/ml Reversed Phase cartages (Part No. 8B-S043-TAK) and pass through it using positive pressure manifold. Subsequently, 50 µL of filtrate was mixed with 50 µL of water. 10µL was injected into the LC-MS/MS system.

Compounds were identified based on retention time and m/z match to injections of authentic standards. Quantification was performed using Waters, Target Lynx™ Application Manage Quantitative software. Peak areas were measured from extracted ion chromatograms of [M-H]-metabolite ions with ± 70 ppm detection windows centered on the theoretical mass. Peak areas from internal standards were measured using an identical procedure; however, values were only used to verify instrument stability and not used in endogenous metabolite quantification.

All standards were reconstituted in (Water and Acetonitrile, 50:50) to prepare 1 mg/ml stock solutions. A standard curve at the range of 5-1000 ng/ml of each analyte was produced from stock solutions in the relevant matrix. 100 µl of each standard was mixed to obtain a mixed standard solution to further prepare a seven-point standard calibration curve within a range of 5-1000 ng/ml. 10 µg/ml of each internal standard (TCA-1 and TCA-2) were spiked in to all samples as an internal standard (final concentration) to allow quantification based on the ratio of the internal standard to each intermediate peak (AUC). Waters Acuity-UPLC coupled with TQD Triple Quad LC-MS/MS mass spectrometer (Waters, Milford, US) was used to quantify the above metabolites. Phenomenex (Torrance, CA) Luna-NH2, 2.0 × 150 mm, 3 µm column was used to achieve an optimal separation of all TCA intermediates and to obtain good peak shapes. The flow rate of 0.3 ml/min was used with a mobile phase A (5 mM NH4OAc buffer at pH 9.9) and mobile phase B (Acetonitrile). Liquid chromatography consisted of a HILIC gradient programs used with mobile Phase A & B listed in table 1. Column temperature was optimized at 27 °C for best peak shape. Electrospray ionization (ESI) negative mode (ESI −) was used for the TCA cycle intermediates. 10 µl was used for the injection volume and the auto-sampler was maintained at 4∘C. Mass spectrometry parameters were modified (Yuan M, 2019) and optimized to obtained best sensitivity and listed in Table 2. The LC eluent flow was sprayed into the mass spectrometer interface without splitting. TCA cycle intermediates were monitored by tandem MS using multiple reaction monitoring (MRM) mode with negative polarity. Identification was achieved based on retention time and product ions. Table 2 summarizes the monitored ions MRM and the optimized MS operating parameters of the analytes. The parameters for TQD mass spectrometry equipped with an ESI probe: Capillary: voltage, 3.5 kV for negative mode: Source Temp: 120 °C: Desolvation temp: 450 °C; Cone gas flow: 120 L/Hr: Desolvation gas flow: 800 L/Hr Collision gas flow: 0.20 mL/min and Nebulizer gas flow: 7 Bar. All metabolites quantification were performed with peak area ratios (PARs) of the analyte peak area to the ISTD peak. The calibration curve, prepared in control matrix, was constructed using PARs of the calibration samples by applying a one/concentration weighting (1/x) linear regression model. All QC sample concentrations were then calculated from their PARs against the calibration curve.

### Relative Quantitation of ^13^C6 flux in TCA cycle Citrate

TQD mass spectrometry was operated at ESI negative mode as described in the previous section. The column effluent was monitored by negative ion electrospray (ESI-) using multiple reaction monitoring (MRM). The primary and confirmatory in-silico MRM transitions (Listed in table 3) were created based on the possible precursor and fragment ions and used quantitation to monitor the retention time and variations. Source and compound parameters were optimized. Target Lynx software was used for relative quantitation of all listed metabolites in the samples with (AUC) compared within two different types of cells (Pre OL and OL)

### Flourmyelin stain

20-30um mouse brain sections were cut on a freezing cryostat and allowed to air dry for 15min. The tissues were rehydrated in P.B.S for another 10min. before applying the Flourmyelin Flourescent dye (Thermofisher: F34651; 75uL each section, concentration; 1:300) conjugated to the FITC. The dye reaction was allowed occur for 15 min, at room temperature in dark and the sections were extensively washed with P.B.S for 3 times and counter stained for DAPI. The sections were further air dehydrated and mounted in flour shield medium. The dry settled slides were imaged using laser scanning confocal microscopy.

### Immune labelling of tissues, cells and imaging methods

All the immune labeling for the cells and tissues was performed as per our already described methods [41]. The antibodies for imaging studies used were: O4: RND Systems-MAB1326, MBP: Cell Signaling 78896, Flag:, Pdhk1: Catalog: Cell Signaling 3820, p-Pdh Ser-293:, For the quantification mature oligodendrocytes, the MBP expressing OLs were collected for the scrambled vector and *h*Pdhk1 transfected slides followed as random snapshots from the laser scanning confocal microscope set to aperture of 1 airy units in the acquisition mode. Multiple snapshots were recorded from both the groups and number of MBP+ OLs were quantified suing Image-J software. The representative pictures from both the groups are displayed. For the Olig1 nuclear to the cytoplasmic shutting studies, the laser scanning confocal microscope was set to acquisition at 1 airy units and multiple snapshots of the Olig1 expressing Pre-OLs were captured. The cytoplasmic expression of the Olig1 in all the groups was determined by the Imaje-J analysis of 15-20 cells in the each group.

### Immunoblotting and Co-Immunoprecipitation (Co-IP)

Immunoblotting was conducted as per well-established methods shown elsewhere [41].Antibodies used are Pdhk2: SC517284, Pdhk3: SC365378, Pdhk4: SC51861, Pdh total: For Olig1 immunoprecipitation reactions, we used the Olig1 antibody (Catalog: SC166264). The cell and tissue lysates were prepared in the protein lysis buffer and incubated with either Olig1 antibody (1:100) or isotype matched IgG antibody (1:100) for 15-20 min on the ice followed by overnight rotation at 4C. After the lysates were processed as per our already described method of immunoprecipitation [41]. The precipitates were further probed for acetyl modifications suing the acetyl-lysine antibody with appropriate inputs and loading controls.

### Pdh Activity determination

The Pdh activity in the cells or tissues lysates was determined using the Sigma Pdh activity colorimetric kit (Catalog: MAK183) as per the manufacturer’s instructions. The activity is normalized to the 5uL of the protein lysate.

### Cuprizone model of demyelination and remyelination

All the animal experiments mentioned in this article were performed as per the IACUC policies in line with the local and federal requirements with animal welfare assurance number D16-00090.The mice were caged n=6/cage with free access to food and water ad libitum with a 12:12 light and dark cycle. 8-10 weeks old C57BL6/J mice of mixed genders were allotted to the groups. Group-1 (n=8): No treatment, Group-2 (n=20): Diet was changed with the Cuprizone diet (0.25% in the chow, Envigo U.S.A). The diet was monitored every alternate day. On average the diet was changed at 4 days interval. The mice from the cuprizone diet were sacrificed at 1 week post introduction, 4 weeks after introduction. Some of the mice (n=5) were taken off the cuprizone diet at 4 weeks post introduction and given normal diet for another 3 weeks and then sacrificed with the normal control group-1. Mice were euthanized using 3.5% CO_2_ asphyxiation and followed by the secondary method of euthanasia either suing cervical dislocation or needle induced pneumothorax. For western blotting the tissues were immediately snap frozen. For imaging purposes, the mice were perfused with 0.95% saline and then 4% paraformaldehyde. The tissues were then again stored in the paraformaldehyde solution for additional 4-5 days.

## Statistics

All the data presented in the manuscript is Mean ± S.E.M. The power of the experiments in terms of the “n” is described in the figure legends. The student’s t-tests were employed to discern differences between the two groups for 1 variable. For more than 2 two groups the differences were determined by using an ordinary ANOVA test. To compare, the multiple groups the post hoc analysis was conducted using Tuckey’s test. Differences between groups ≤ 0.05 was considered to be significant with the given sample size in a particular experiment.

## Discussion

Our findings highlight a unique role of integrated signal and cellular metabolism program via Pdh which regulates the maturation of the pre-OLs into the myelinating OLs. These findings are especially significant in the context of demyelinating diseases like MS, which has been reported to block the terminal maturation of pre-OLs into the OLs [42-44]. With the current literature demonstrating growing consensus that MS inflammation affects oligodendrocytes development adversely [45], our studies emphasize a key role of Pdh in OL maturation. Any alterations in the Pdh activity or expression during MS can negatively impact the myelination by halting the terminal maturation of the Pre-OLs.

Previous studies have clearly demonstrated the significance of mitochondrial respiration in oligodendrocyte maturation [24] [46]. However, the data developed previously focused more on the OPC and OLs [24] and did not address the intermediate and the immediate stage of pre-OLs prior to the maturation into the OLs. Our work progresses the existing idea to an important transition from Pre-OLs to the OLs. These findings are directly relevant the human disease given MS lesions show significant number of pre-OLs which possibly fail to mature into the OLs [19]. Hence during MS, the OPCs may differentiate into the pre-OLs but fail to mature into myelinating OLs. In an another landmark study using radioactive carbon dating studies in World War II MS patients, the investigators did not find any defect in the rates of the OPC proliferation [47]. But they didn’t find new mature OLs in the MS lesions thus highlighting the bloc of the maturation. Collectively, these show that demyelination leads to the OPC proliferation but proliferation doesn’t translate into the maturation of the OLs thus impacting the remyelination. In line with these, our early work in the mouse model of the M.S has shown shortening of the cell cycle of the OPC progenitors in the preclinical models of the M.S[48]. Our current work further shows that Pdh activation during Pre-OL maturation into the OLs. Hence Pdh alterations in MS can impact the maturation of the Pre-OLs without affecting the proliferation rates and thus halting the remyelination process and critically contribute to the overall demyelination and injury and apart from contributing to the disability.

There has been a scarcity of investigations at the signal transduction level attempting to understand the signal biology underpinning the shifting metabolic demand during oligodendrocyte development. Apart from increasing oxygen consumption, our metabolic profile studies clearly reveal increased acetyl-CoA in mature oligodendrocytes. These findings suggested that Pdh may have a function in relaying pyruvate from glycolysis to acetyl-CoA, allowing for increased oxygen consumption and, as a result, *de-novo* synthesis of fatty acids, both of which play important roles in myelin biology [49-54]. Our findings show only pdhk1 is inhibited during the conversion of the pre-OLs into oligodendrocytes. We also saw a trend in the rise of the Pdhk4 during this transition which in theory should deactivate Pdh. Given we find a clear rise in the Pdh activity during Pre-OL transition into the OLs, the rise in Pdhk4 may not be significant. Nevertheless, these data underline the importance of context dependent modulation of a specific kinases achieving terminal maturation in the oligodendrocytes. Although our findings are unique, given the variety of methods used to differentiate OPCs into oligodendrocytes, it is possible that other kinase isoform inhibition patterns will be uncovered in future. Grossly pyruvate dehydrogenase complex deficiency leads to the neurodegeneration and underdeveloped grey and white matter in the humans [55]. When pyruvate dehydrogenase complex (PDC) was conditionally deleted in oligodendrocytes, it had no significant effect on myelination [56]. Our findings reveal that blocking Pdh activity via overexpression of Pdhk1 causes Pre-OLs to fail to mature into mature OLs. These findings suggest that Pdh activity via Pdhk1 may be important in the terminal maturation of the Pre-OLs. Once the oligodendrocytes reach maturity, they may no longer be entirely reliant on the oxidation of pyruvate into acetyl-CoA via its oxidative decarboxylation through the Pdh. This is evidenced by the unaffected myelin after the conditional loss of pyruvate dehydrogenase complex in the mature oligodendrocytes [56]. It’s possible that once the OLs mature, they might have developed alterative metabolic processing to acetyl-coA. This is supported by the fact that mature oligodendrocytes utilize 10% pyruvate via pyruvate carboxylation [25] thus reducing the need for Pdh dependent acetyl-CoA generation. When the preclinical M.S model mice are treated with acetate, a direct precursor of acetyl-CoA, the mice showed reduction in the disability and better myelination outcomes [57]. Taken together these in conjunction with our data point to need for Pdh dependent acetyl-CoA generation during the Pre-OL maturation might be critical for the remyelination and reduction of disease burden.

The unifying mechanisms of how signal-mediated alterations in metabolic rewiring affect the cellular milieu are still in their infancy. Recent studies have focused on the role of the basic helix loop helix transcription factor Olig1 in oligodendrocytes differentiation and maturation [58-61]. Numerous studies have revealed that Olig1 expression is necessary for the fate attainment of OPCs to oligodendrocytes [58,62,36,63]. It has been reported that Olig1 modification at different residues affects OL maturation from the Pre-OLs. Acetylation of Olig1 was reported to regulate oligodendrocyte maturation through Olig1 nuclear to the cytoplasmic shuttling allowing for massive membrane expansion, which is a prerequisite for myelination [36]. Our work confirms this finding and we also see less nuclear sequestration and higher cytoplasmic expression of Olig1 during the Pre-OL maturation into the OLs. This transition was however blocked by the Pdhk1 mediated Pdh deactivation. These results show a previously unknown metabolic alteration controlling the OL maturation. Thus inhibiting Pdhk1 would allow an abundant acetyl-CoA pool, increasing the likelihood of Olig1-acetylation. Previous studies using *in vivo* using ^13^C-tracing *in-vivo* found increased lactate specific to higher demyelination in the brains of cuprizone model of demyelination [39] which can be a direct outcome of the Pdh deactivation. Indeed, authors in the latter study identified an increased Pdhk1 in numerous cell types during cuprizone demyelination. However, Pdhk1 expression in pre-OLs was not investigated. Our results corresponds with this study in that during cuprizone-induced demyelination, Pdh activity is reduced during the demyelination phase and returns during the remyelination. In line with these, the pre-OLs express higher levels of Pdhk1 in demyelination phase compared to baseline and these changes were reversed after cuprizone was removed. This notion was further strengthened by the fact that the Olig1-acetylation was repressed during the demyelination phase and restored when cuprizone was removed from diet for 3-weeks. In line with previous reports, we discovered that while the baseline exhibits modest cytoplasmic localization of Olig1, the demyelination phase demonstrates higher sequestration of Olig1 in the nucleus of the premature oligodendrocytes. Cuprizone withdrawal, on the other hand, resulted in a greater cytoplasmic localization of Olig1 in premature oligodendrocytes, allowing them to mature faster.

Concluding our work progresses the oligodendrocyte biology and highlights inter wiring of the signal transduction and cellular metabolism in the oligodendrocyte physiology and maturation with direct implications on the demyelinating diseases of CNS.

## Statements and declarations

### 1. Ethics Approval

These studies were performed after due approval from the ICUC committee from Henry Ford Health System (complying with the national guidelines for the use of animal subjects in the research) vide number protocol number 1363 dated June 01, 2022. The biological materials was obtained as detailed in the humane endpoints instituted in the study protocols.

### 2. Consent to participate

No human subjects involved in study.

### 3. Consent for Publication

No human subjects involved in study.

## 6. Author Contributions: Contributions

M Sajjad, Faraz Rashid and Insha Zahoor collected the data, M Sajjad, Insha Zahoor and Faraz Rashid analyzed the data. Ramandeep Rattan helped in the analysis of the LC/MS data. Sajjad M wrote the manuscript and Shailendra Giri edited the manuscript. All the authors have approved the final data and the final version of the manuscript.

## 7. Acknowledgements

Authors would like to thank confocal laser scanning facility at the department of neurosurgery, Henry Ford Health (HFH). We would like to thank Polyplus for sharing transfection reagents with us as a generous gift.

## 8. Availability of data and materials: Acknowledgements

All the data sets generated are not publically available do to the originality of the work and being submitted first time for the publication. However these data are retained and will be made available after a reasonable request is made to the corresponding authors.

